# Self-organised symmetry breaking in zebrafish reveals feedback from morphogenesis to pattern formation

**DOI:** 10.1101/769257

**Authors:** Vikas Trivedi, Timothy Fulton, Andrea Attardi, Kerim Anlas, Chaitanya Dingare, Alfonso Martinez-Arias, Benjamin Steventon

**Affiliations:** Department of Genetics, University of Cambridge, Cambridge, CB2 3EH, UK; European Molecular Biology Laboratories (EMBL), Barcelona, 08003, Spain; Biotechnology for Industry and Scientific Research (BIRS), University of Palermo, Palermo, Italy; Max Planck Institute of Molecular Cell Biology and Genetics, Dresden, Germany

## Abstract

A fundamental question in developmental biology is how the early embryo breaks initial symmetry to establish the spatial coordinate system later important for the organisation of the embryonic body plan. In zebrafish, this is thought to depend on the inheritance of maternal mRNAs [1–3], cortical rotation to generate a dorsal pole of beta-catenin activity [4–8] and the release of Nodal signals from the yolk syncytial layer (YSL) [9–12]. Recent work aggregating mouse embryonic stem cells has shown that symmetry breaking can occur in the absence of extra-embryonic tissue [19,20]. To test whether this is also true in zebrafish, we separated embryonic cells from the yolk and allowed them to develop as aggregates. These aggregates break symmetry autonomously to form elongated structures with an anterior-posterior pattern. Extensive cell mixing shows that any pre-existing asymmetry is lost prior to the breaking morphological symmetry, revealing that the maternal pre-pattern is not strictly required for early embryo patterning. Following early signalling events after isolation of embryonic cells reveals that a pole of Nodal activity precedes and is required for elongation. The blocking of PCP-dependent convergence and extension movements disrupts the establishment of opposing poles of BMP and Wnt/TCF activity and the patterning of anterior-posterior neural tissue. These results lead us to suggest that convergence and extension plays a causal role in the establishment of morphogen gradients and pattern formation during zebrafish gastrulation.

---

Our current understanding of pattern formation during early development relies heavily on the notion of opposing signalling gradients that set-up rudimentary body plans [17]. These gradients establish cell fates in space that in turn lead to the population specific cell behaviours that dictate the complex cell and tissue rearrangement of gastrulation and axial elongation. In zebrafish, opposing Nodal and BMP signalling gradients are thought to be necessary and sufficient for the establishment of the body plan as shown by experiments in which deployment of such gradients in animal caps leads to the formation of a complete AP axis [13]. In addition to controlling cell fate assignments, a recent study has demonstrated that Nodal signalling is a key driver of convergence and extension movements and is sufficient to generate these behaviours when expressed within zebrafish animal caps [14]. Furthermore, BMP levels have been shown to be important for controlling cell movements during both gastrulation [21] and posterior body elongation [22]. These observations raise the possibility that opposing BMP and nodal signalling gradients are upstream of both morphogenesis and patterning. However, the causal relationships of these processes are difficult to dissociate in situations where continuous external signaling sources are present, either from overexpression experiments or from the extra-embryonic signals present during early development. To follow how cells can develop and pattern in the absence of external signals, we used primary culture of cells from zebrafish embryos at the 256 cell stage i.e. before the midblastula transition (Figure 1A). This stage also precedes the formation of the zebrafish extra-embryonic yolk syncytial YSL which has been suggested to shape the embryonic axes through mesendodermal induction [9–12] and regulation of epiboly [15].

**Fig. 1.**
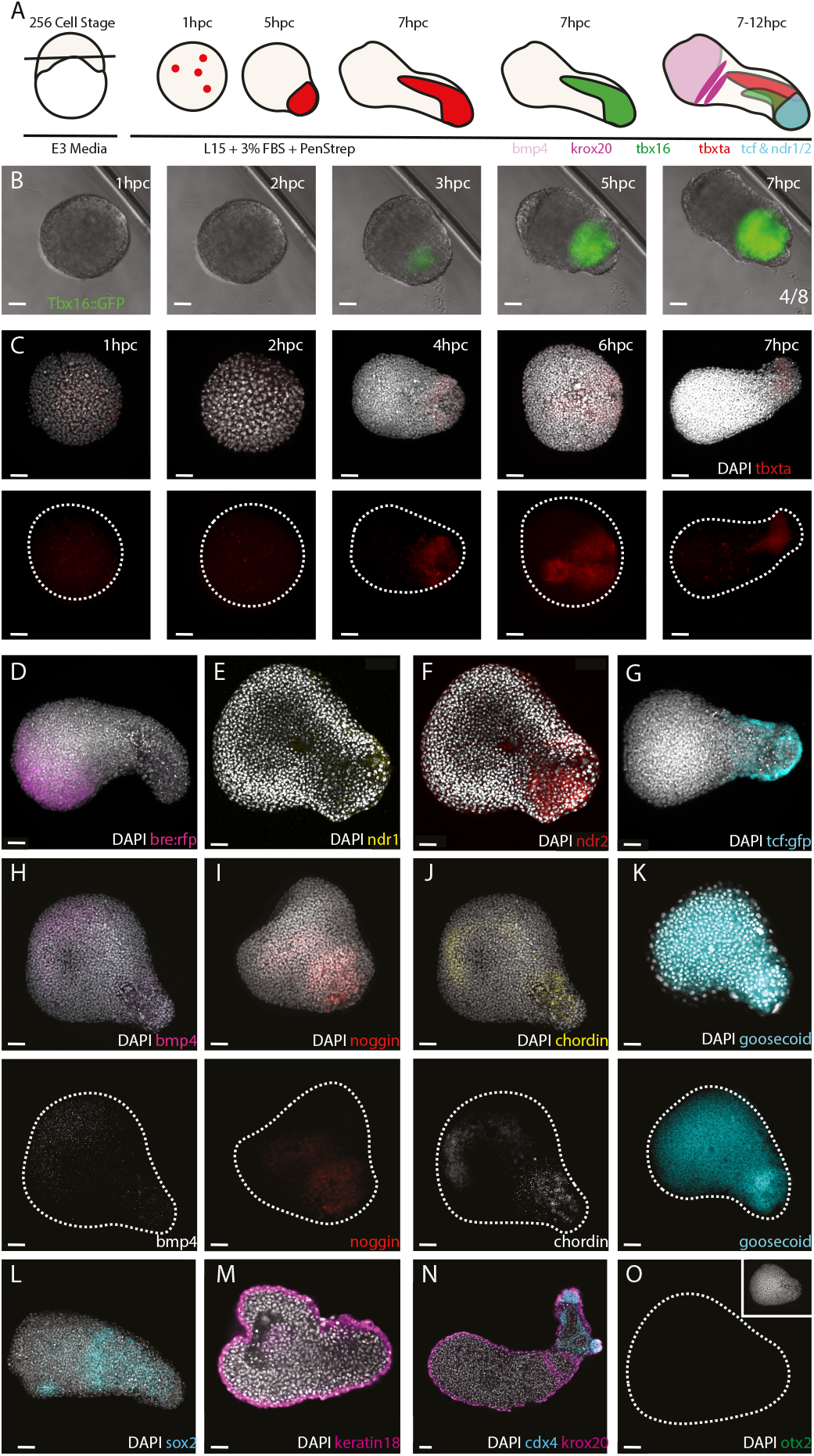
Symmetry breaking and axial patterning can occur in the absence of extra-embryonic signals. (A) Explants of early embryonic blastomeres taken at the 256 cell stage demonstrate elongation and mesendomeral induction visualised through expression of a (B) Tbx16:GFP reporter (n=4/8) and (C) tbxta mRNA (n= 1hpc 4/4; 2hpc 5/6; 3hpc 5/6; 5hpc 4/11; 7hpc 6/6). At the opposite pole to tbxta expression, (D) bmp4 activity is observed (n=4/6) (E-F) alongside the tbxta pole, nodal ligands ndr1 and ndr2 are expressed (n=5/6). (G) TCF/beta catenin activity occurs alongside with the tbxta pole (n=7/8). The region of BMP activity correlated with (H) expression of the BMP ligand bmp4 (n=8/9). The high Nodal/Wnt domain also expresses organiser marker genes including: (I) noggin (n=3/3), (J) chordin (n=6/6) and (K) goosecoid (n=2/4). The remaining tissue expresses patterned neural markers including (L) sox2 (n=3/6) and (N) cdx4 (n=5/5) and krox20 (n=7/8). (M) Krt18, an epidermal marker, is expressed in the outermost layer (n=4/4). (O) Expression of otx2 is not observed in the explants (n=4/4). Scale = 50*μ*m. n=Expression observed/total number imaged.

Explants from embryos at different stages between the 64 cell and 512 cells, all exhibited a similar behaviour (Figure S1A-B): they self-organised to form polarised aggregates with a protrusion emerging from one pole. We focused our studies on 256 cells stage embryos that exhibit this behaviour in more than 60% of explants from each experiment (n=20, 80-100 explants per experiment; Video S1). Quantification of aspect ratio of the longest vs. shortest axis of each aggregate over time revealed a coordinated onset of elongation at 7 hours post culture (hpc; Figure S1A), demonstrating a degree of synchrony in the symmetry breaking event. Explants from Tbx16:GFP reporter embryos [16] revealed mesoderm specification within the elongating end of the aggregate (Figure 1B; Video S2), accompanied by polarised expression of tbxta (Figure 1C). These results showed how symmetry breaking and mesoderm patterning can occur spontaneously in embryonic cells when separated from the yolk. Fixing aggregates at the onset of elongation revealed a pole of BMP activity (Figure 1D) opposite the elongating end that was characterized by ndr1 and ndr2 expression and signalling as revealed by a phospho-Smad2/3 expression pole (Figure 1E,F and Figure S1C). In addition, we observed a gradient of Wnt/beta-catenin activity extending from the most posterior end of the aggregate (Figure 1G, Figure S1D and Video S3). Opposite the end expressing bmp4 (Figure 1H) is a region expressing noggin (Figure 1I) chordin (Figure 1J) and goosecoid (Figure 1K), that was likely responsible for the absence of BMP activity at this pole. As in the embryo, the domains of high Wnt activity and ndr1/2 expression correlated with the overlapping expression of these organiser genes. The observed poles of signalling activity translated into an approximate anterior-patterning of neural tissue at 12hpc (as judged by sox2 expression; Fig 1L), with cdx4 expression in the posterior region and krox20 at an intermediate position (Figure 1N). Surrounding this we notice a layer of krt18 positive epidermal tissue (Figure 1M). However, we see no evidence for the existence of forebrain markers, as shown here for otx2 (Figure 1O). This suggests that additional signal modulation might be required to fully pattern the neural tissue observed in early embryo aggregates. Our results demonstrate that opposing poles of signalling activity can form in these aggregates, establish a gastrula organiser and pattern a rudimentary anterior-posterior neural axis in the absence of continued signals from the yolk and the YSL. The observed axial emergence in the absence of any inductive signals from the YSL is also supported by previous works on the dispensability of extra-embryonic YSL in mesendodermal induction [18]. In reference to the self-organisation of mammalian cells into polarized ‘gastruloids’ [19,20], we refer to the self-organised aggregates of early embryonic zebrafish cells as *‘pescoids’*.

In the embryo mesoderm is specified in part by the inheritance of maternal mRNAs in the vegetal-most blastomeres that remain in direct contact with the yolk through to the 128-cell stage [1–3] and might be important for setting up an initial symmetry breaking event that leads to the emergence of the opposing signalling gradients described above. To test whether pre-existing spatial asymmetries are required for continued polarised tbxta expression, we dissociated explanted cells and reaggregated them to determine whether polarization is observed upon removal of any pre-existing asymmetry in the re-aggregates (Figure 2A). In many cases, reaggregation was not complete, leading to the formation of smaller pescoids. However, in 7/10 of these reduced-sized pescoids, a polarized tip of tbxta expression was still observed (Figure S2A). When most cells were reaggregated, a clear elongated morphology was observed together with tbxta polarization (Figure 2B) as in non-dissociated pescoids (Figure 1C).

**Fig. 2.**
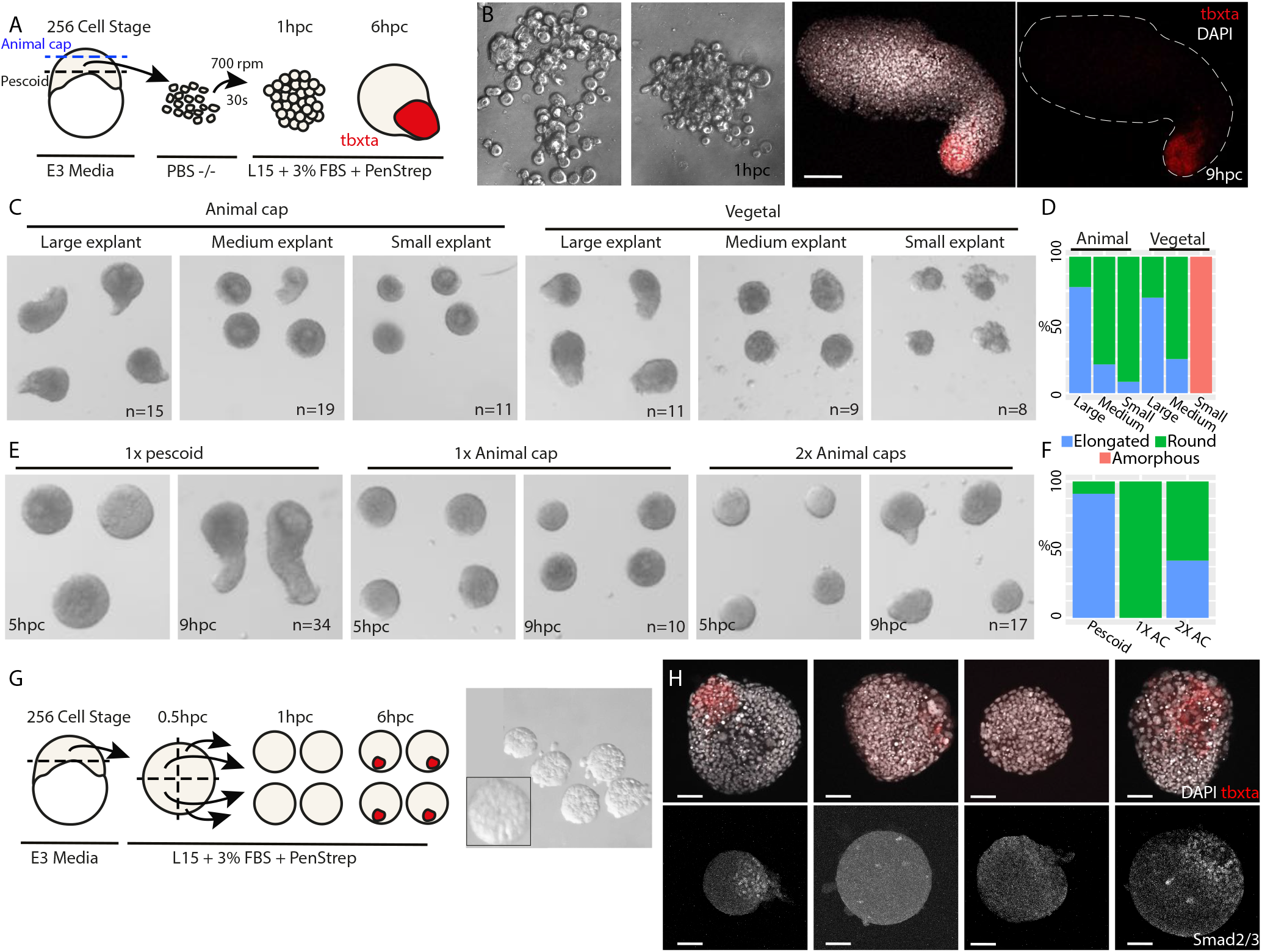
Symmetry breaking is robust to experimental cell mixing. (A) Disassociation and reaggregation of explanted cells results in polarisation of tbxta expression (n=8/8; Tbxta expressed/total imaged) and (B) infrequently, in elongation of the aggregate (n=2 observed elongating). (C-D) Neither animal nor vegetal explants demonstrate a bias towards elongation however the size of the tissue correlates with elongation potential (n=8 expants minimum per class). (E-F) Full pescoids demonstrate a clear and robust elongation however this is not wholly reproduced by aggregation of two animal caps together (n= 5/12 elongated/total) and was never seen in single animal cap explants (n=0/10 elongated/total). (G) Quartering of single pescoids also reduced the potential of pescoids to elongate however (H Upper) tbxta expression was observed (n=26/32 expression observed/total imaged) in the (H Lower) absence of robust Smad2/3 activity (n=6/21 activity observed/total imaged). Scale = (B) 50*μ*m; (H) 30*μ*m.

If the inheritance of maternal mRNAs were important for the establishment of symmetry breaking in pescoids, we would expect that vegetal explants would have a higher capacity to elongate than animal ones. When taking both animal and vegetal (Figure 2C) regions at a range of sizes, we found no bias towards vegetal explants in their ability to elongate although tissue size was a key factor (Figure 2D). Indeed, complete pescoids elongated to a much greater extent than either poles alone suggesting that a complete set of cells was important for pescoid elongation (Figure 2E). To test whether this size dependency might explain the absence of elongation in animal caps, we dissociated and reaggregated 2 animal caps together. In some cases, this led to the formation of a protrusion, however this was again dependent on aggregate size. In no cases were protrusions observed in single reaggregated animal caps (Figure 2E,F). This size dependency was also confirmed when cutting un-dissociated pescoids into smaller portions (Figure 2G). Pescoids cut into halves are able to elongate (Figure S2B) while individual quarters are able to express tbxta but do not elongate (Figure 2H, upper row; Figure S2C). Of the quarter-sized pescoids, we observed a pole of Smad2/3 activity in only a few cases (Figure 2I, lower row; Figure S2D). Taken together, these results suggest that aggregate size is the major determining factor for early embryonic cells to break symmetry and elongate, rather than the inheritance of a pre-pattern from maternal deposited mRNAs.

The ability of pescoids to undergo symmetry breaking and elongation in the absence of a pre-pattern, led us to question how cells behave in early pescoids. We used light sheet imaging (SPIM) of pescoids immediately after harvesting. Cells undergo several rounds of rapid division and increase by about 20-fold over a period of 2.5 hrs (Figure 3A, B, Video S4). In order to understand the nature of division patterns, we estimated the increase in the number of cells in two scenarios: (i) if all cells were to divide synchronously every 20 min (Fig 3B, blue graph) or (ii) only a random sub-population (≤ 50%) of cells were to divide every 20 min (Figure 3B, orange graph). Compared to these estimates it is clear that there is a degree of asynchrony in the rate of divisions across the early pescoids. Analysis of the direction of division, further showed that there is no spatial pattern of divisions (Figure 3C) and as a result of which prior lineage-based pre-patterns would become homogenized owing to these rapid, asynchronous divisions within the pescoids before elongation. To determine where cell movements in the pescoids might also contribute to the erasure of any pre-existing spatial pre-pattern, we performed small photo-labels of cells at one edge of the pescoid and observed their distribution 3 hours later (Figure 3D). In all case, a complete mixing of labelled and unlabelled cells was observed (Figure 3D, Figure S3). These results reveal that pescoids undergo extensive cell mixing at early stages that effectively remove any pre-pattern that could be produced from the inheritance of maternal mRNAs at the vegetal pole. Accordingly, HCR staining of animal caps revealed the expression of the maternally deposited mRNA eomesodermin, as well as the early mesodermal markers tbxta and goosecoid (Figure 3E). In addition, time-lapse movies of animal caps showed a clear expression of the Tbx16:GFP reporter (Video S5). Together, these results suggested to us that maternal inheritance of mesodermal specification is inherited to both vegetal and animal blastomeres by the 256-cell stage However, additional size-dependent signalling events are required to generate pescoid elongation and neural patterning.

**Fig. 3.**
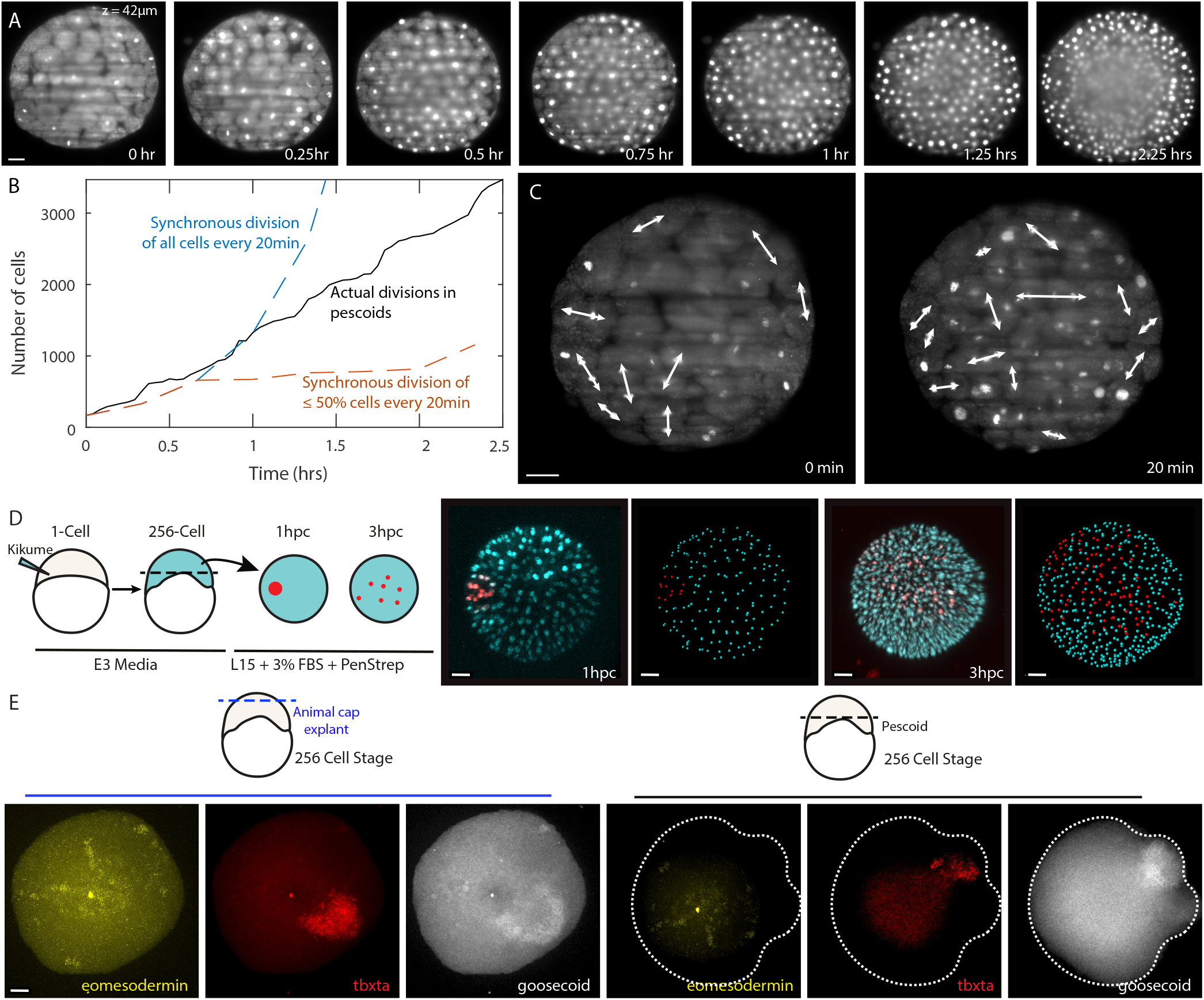
Lineage and spatial pre-patterns are lost due to extensive cell divisions and cell mixing. (A) Cells in pescoid undergo rapid cell divisions as seen in images acquired on SPIM. The data shown is for an optical section 42um below the surface. (B) Number of cells in pescoid counted based on image segmentation (black curve). Dashed curves are estimates of the number of cells, starting from the number of spots segmented at t = 0, if all cells divided synchronously every 20 min (blue line) or only a random sub-population (*leq*50%) of cells divided every 20 min (orange line). (C) Cells in pescoid divide randomly with no preference for direction of division, leading to mixing of cells. (D) Mixing of labelled and unlabelled cells is observed in pescoids injected 3 hrs prior with kikume and then a small population of cells is photo converted for labelling (n=6 replicates, all demonstrating cell mixing). (E) HCR staining of animal cap explants and pescoids reveals a similar expression of eomesodermin (pescoids 4/4; animal caps 6/6), tbxta (pescoids 5/6; animal caps 3/4) and goosecoid (pescoids 3/6; animal caps 2/4) in both animal caps and full pescoids suggesting that animal caps are able to break symmetry but require a size-dependent signalling event to achieve elongation and neural patterning. n=expression observed/total number imaged in (E).

Having established that pescoids can break initial symmetry, establish opposing signalling poles, and pattern anterior-posterior neural axis, we next sought to follow the temporal order of these events. To determine the earliest signalling event associated with elongation, we fixed pescoids at intermediate stages between initial culture and protrusion formation at 5hpc and stained for diphosphorylated ERK-1&2 (Figure S4A), beta-catenin (Figure S4B) and phospho-Smad 2/3 (Figure 4A). The earliest signal to associate with the elongated pole and tbxta expression (Figure 1C), is Smad2/3 signalling (Figure 4A), suggesting that a self-organised pole of Nodal activity is one of the earliest steps driving elongation. Furthermore, blocking Nodal receptor activity with SB505124 inhibits pescoid elongation (Figure 4B). This is in line with recent work showing that Nodal signalling is important to drive convergence extension movements within animal caps [23].

**Fig. 4.**
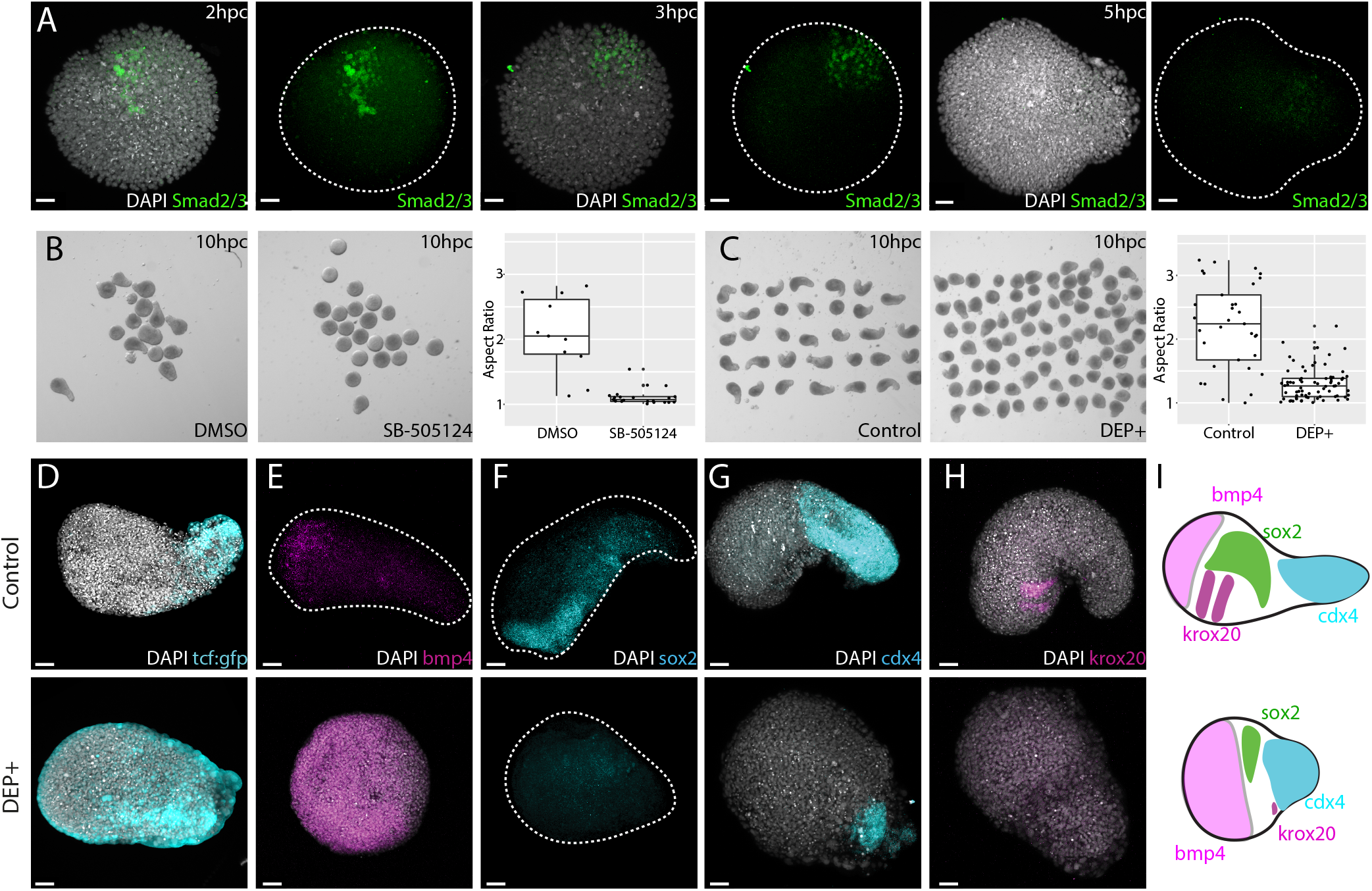
Nodal dependent convergence and extension movements lie upstream of gradient formation and anterior-posterior patterning.. (A) The presence of phospho-Smad 2/3 was assayed across a time course revealing smad2/3 correlates with the onset of tbxta expression (smad positive n= 2hpc 3/10; 3hpc 2/10; 5hpc 4/11). (B) Using SB505124, nodal activity was inhibited for the first two hours of culture prior to being washed away. This early treatment is sufficient to block the elongation of the pescoid at 10hpc. (C) Injection of a dominant negative Dishevelled (dsh-DEP+) also inhibits pescoid elongation through blocking of the PCP pathway. The absence of elongation, (D) the Wnt/beta-catenin domain is no longer located only at the tbxta positive pole as in controls (11/13 show expansion of TCF:GFP away from the tbxta pole) and (E) an expansion of the BMP4 domain occurs (7/12 show expansion of bmp4). This results in a smaller (F) sox2 (7/9 show smaller sox2 expression domains) and (G) expanded cdx4 domain (4/6 have expanded cdx4 expression). (H) Krox20 expression is normally observed in two clearly defined stripes however in DEP+ injected pescoids, this expression pattern was lost (5/6 show no clear two stripe pattern of krox20). (L) These results together demonstrate that opposing BMP/Wnt gradients, which give rise to the anterior posterior pattern of the neural tissue, lie downstream of Nodal and PCP dependent convergence and extension. Scale = 50*μ*m.

As Nodal activation and convergence and extension precede the establishment of BMP and Wnt signalling poles and anterior-posterior patterning, we hypothesised that elongation may be important for establishing embryonic patterning by these signals, as has been previously shown for retinoic acid [28]. To test this, we specifically inhibited the PCP pathway using a dominant negative version of Dishevelled (Dsh− DEP+) that in Xenopus inhibits activin induced axial elongation, without perturbing Wnt/beta-catenin activity [14]. This blocked pescoid elongation (Figure 4C) while still allowing for a pole of Smad2/3 activity to form (Figure 4D). In this context, Wnt/beta-catenin signalling is no longer localized to the tbxta positive pole as compared to controls (Figure 4D, Figure S4C). Under these conditions, most of the aggregate expressed bmp4 (Figure 4E). Concordantly, these pescoids had a smaller domain of sox2 positive tissue (Figure 4F) that was positive for the posterior marker cdx4 (Figure 4G) and the hindbrain marker krox20 was either reduced or completely absent (Figure 4H). In no cases did we observe the double stripes of expression seen in controls. Taken together, these results showed that Nodal and PCP dependent convergence extension movements lie upstream of the establishment of opposing BMP/Wnt gradients and the anterior-posterior patterning of neural tissue (Figure 4I).

Early explants of embryonic cells allow a test of their potential without extraembryonic tissues. When such explants are and allowed to aggregate and develop in culture, they reveal a previously unnoticed feedback mechanism between morphogenesis and cell fate assignment. We find that convergence and extension movements are important for establishing appropriate distance from an anterior source of BMP and a posterior source of Wnt/beta-catenin activity. This enables a significant proportion of tissue to be specified as hindbrain (marked by krox20) in a region that is low in BMP and moderate Wnt/beta-catenin activity. Posterior to this lies a high Wnt/beta-catenin, low BMP domain expresses markers of posterior neural (cdx4) and mesodermal expression (ntla, tbx16). Importantly however, we never see otx2 expression, suggesting that additional spatial separation is required to create a region that is both low in Wnt/beta-catenin and BMP to specify forebrain. During normal gastrulation movements, this occurs as the prechordal mesoderm moves anteriorly and continues to inhibit both BMP and Wnt activity [30]. Whether the absence of forebrain specification in pescoids is due to the lack of additional extra-embryonic signals, or due to the fact that additional morphogenetic events to separate organiser derived signals is an open question.

We observe that mesoderm specification can occur in the absence of a maternal pre-pattern or continued source of Nodal signalling from the YSL. How cells break symmetry to establish a pole of Smad2/3 activity on one side of the aggregate is unknown. Previous studies have shown that a transport and diffusion of signals from the YSL are important for ensuring a high level precision in mesoderm specification [24]. How the diffusion of Ndr1 and Ndr2, and their inhibitor Lefty interact in the context of extensive cell mixing will be essential to obtain a complete picture of how symmetry breaking occurs in both whole embryos and explants. During normal development, it is likely that self-organised symmetry breaking and YSL signal release act together to ensure that embryo patterning is both robust to alterations in external environment and the initial conditions of the fertilized egg.

While the de novo patterning observed in pescoids recapitulates only a portion of the pattern observed during normal embryonic development, this nonetheless reveals the existence of a self-organizing process that might underpin the organization of the primary body axes in a range of species. For example, when mouse embryonic stem cells (mESCs) are allowed to aggregate in 3D to form a spheres (gastruloids) in a medium that lacks spatially localized signalling cues, they display symmetry breaking events in the absence of extraembryonic material that resemble those that we have described here [19,20]. Similar emergence of embryonic pattern has been observed in dissociated and re-aggregated cells from other metazoan species such as hydra [25], Xenopus [26,27] and occurs naturally in Killifish [29]. How each of these examples differ in the precise mechanisms of symmetry breaking and patterning are likely to reveal further insight into how morphogenesis, morphogens and gene-regulatory networks interact to generate pattern during complex morphogenesis.

## Acknowledgments

We would like to thank Carolina Monck and Rohan Sanghera for helping with some experiments and the quantification of pescoid elongation respectively. Many thanks to Caroline Hill for sharing the Nodal and Wnt reporter lines, Michael Lardelli and Simon Wells for sharing the Tbx16::GFP reporter line and the Steven Wilson lab for sharing the TCF::GFP reporter zebrafish. We also thank Gopi Shah at the Mesoscopic Imaging Facility at EMBL Barcelona for help with SPIM imaging

## Author contributions

Conceptualization: V.T, and B.S.; Funding acquisition: V.T., A.M.A. and B.S.; Investigation: T.F., V.T., A.A., K.A., and C.D; Methodology: V.T., A.A. B.S and T.F.; Project administration: V.T. and B.S; Resources: B.S.; Supervision: V.T and B.S.; Validation: T.F. and K.A.; Visualization: V.T. and T.F.; Writing - original draft: V.T. and B.S.; Writing - review and editing: A.M.A, T.F., V.T., and B.S.

