## Supplementary methods and figures for "Self-organised symmetry breaking in zebrafish reveals feedback from morphogenesis to pattern formation" (from another version of this preprint: version 1, 2019-09-14)

**2 Supplementary Figures** **4**

**3 Supplementary Movies** **11**

[1]Department of Genetics, University of Cambridge, Cambridge, CB2 3EH, UK. [2] Current address: European Molecular Biology Laboratories (EMBL), Barcelona, 08003, Spain [3] Biotechnology for Industry and Scientific Research (BIRS), University of Palermo, Palermo, Italy [4]Current address: Max Planck Institute of Molecular Cell Biology and Genetics, Dresden, Germany. [#]Equal contribution. [*]

### 1 Materials and methods

#### 1.1 Zebrafish lines

Adult fish were raised and maintained as described in *A guide for the laboratory use of zebrafish Danio* (Brachydanio) rerio* by Monte Westerfield (1994) in strict accordance with the recommendations of the Animals (Scientific Procedures) Act 1986 Amendment Regulations 2012 following ethical review by the University of Cambridge Animal Welfare and Ethical Review Body (AWERB). Wildtype (WT) embryos were obtained from TL strains. Upon crossing appropriate adult lines, the embryos obtained were raised in E3 media (287.1 mg/L $NaCl$, 13.2 mg/L $KCl$, 47.85 mg/L $CaCl_2.2H_2O$, 80.685 mg/L $MgCl_2\Delta6H_2O$ and 100 $\mu$L of 1% methylene blue in 1L water) at 28.5°C. Transgenic lines for Nodal and Wnt reporters were kindly provided by Caroline Hill lab, TCF::GFP reporter line was provided by the Steven Wilson lab and Tbx16::GFP reporter was obtained from Lardelli lab ($Wells\ S.\ et\ al.\ PLOS\ ONE$ (2011) )

#### 1.2 *In vitro* culture of zebrafish explants (*pescoids*)

Zebrafish embryos (injected or otherwise) were allowed to develop till 32 to 64-cell stage (1.75 - 2 hpf) when they were dechorionated. When the embryos reached 256 cell-stage (2.25 - 2.5hpf), majority of cells were cut using the standard eyelash tool typically used for orienting embryos. Immediately after cutting the cell mass was transferred to Leibovitz's L-15 Medium (ThermoFisher Cat. # 11415049) with 3 $\mu M$ Pen-Strep (Sigma Cat. # P4333-100ML) and with 3%FBS (Biosera Cat. # FB-1090/500) ,where it formed a sphere.

#### 1.3 mRNA injection for photolabelling

Embryos, obtained by crossing WT TL zebrafish adults, were injected with 200pg of nuclear-targeted Kikume mRNA (a kind gift from Ben Martin, Stony Brook University, NY, USA) at one cell stage. Embryos were then raised, at 28.5degC and used for making pescoids at 128 or 256 cell stage. Within 0.5 hour post culture (hpc) the pescoids were embedded in a small drop of 1% methyl cellulose in glass-bottomed dish. About 10-20 nuclei were photo-converted from green to red by rapid exposure to 405nm laser at with a 10X objective. The pescoids were then removed and allowed to grow. They were imaged again at 5hpc.

#### 1.4 Imaging and analysis

For live imaging of pescoids samples were placed in an imaging dish with a #1.5 coverglass bottom and were free floating in wells made from agarose to limit movement around the dish. The imaging plate was then placed on wide field epi-fluorescence microscope (either Zeiss AxioObserver or Nikon, with temperature control) and imaged with a long working distance 10x air objective. For fixed embryos/explants, following the staining protocol, sample was placed in an imaging dish with a #1.5 coverglass bottom in CUBIC clearing solution ($Gomez - Gaviro\ M.V..\ et\ al.\ Development$, 2017) and imaged on confocal microscope (Zeiss LSM 700) using 20X objective. For light sheet imaging, Luxendo (Bruker) MuVi SPIM was used and samples were mounted on 1% (w/v) agarose bed and imaged from the sides. Images are displayed without subtracting background unless otherwise stated. Confocal images are displayed without subtracting background and z-stacks are displayed as maximum intensity projections unless otherwise stated. For analysis, any quantification of shape on widefield images was done manually using line drawing tool in FIJI (we thank Rohan Sanghera for help with analysis). Analysis of SPIM images was done using TGMM tracking software ($Amat,\ et\ al.\ Nature\ Methods$, 2014). Bright-field images of pescoids under different conditions (different starting cell number, media composition, etc) were acquired on a stereo microscope. The colored

(RGB) images were then converted to a gray scale image in FIJI and are displayed without any change of contrast. Unless otherwise stated, all such panels in the same figure are acquired and displayed at the same zoom (magnification). ,

##### 1.5 In situ hybridization chain reaction (HCR)

Embryos/explants were fixed at appropriate stage, using freshly prepared 4% (w/v) paraformaldehyde (PFA) solution in PBS, prior to performing in situ hybridization chain reaction (HCR). HCR probes, hairpins and buffers were purchased from Molecular Technologies (molecularinstruments.org). The protocol used was adapted from the one for whole-mount zebrafish embryos provided in Supplementary Section S7 of *Choi H.M. et al. Development* (2016) and Supplementary Section S2 of *Choi H.M. et al. Development* (2018). Hairpin sequences for the different systems were same as those described in *Choi H.M. et al. Development* (2016).

#### 2 Supplementary Figures

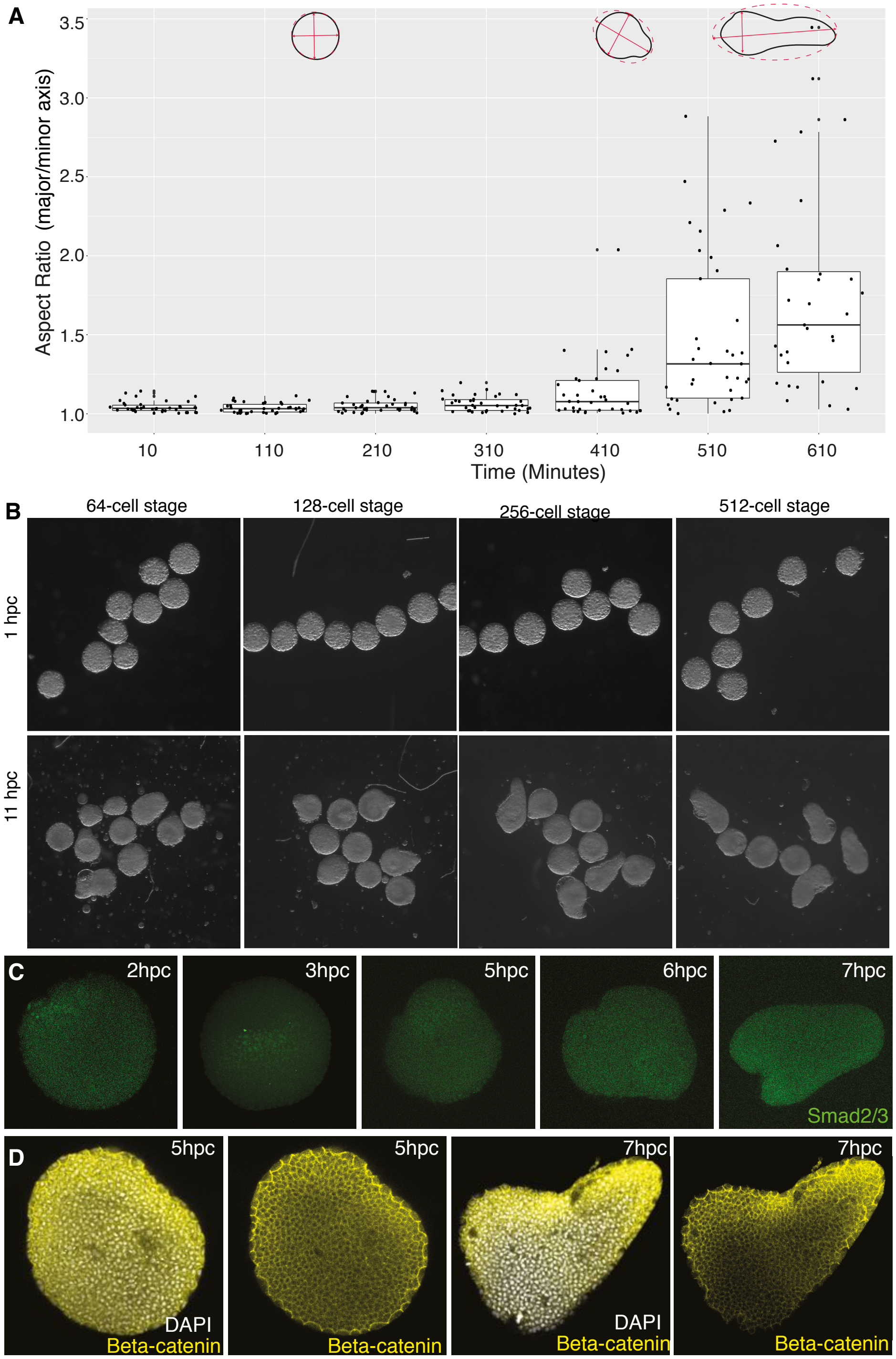

**Figure S1.** Elongation of pescoids

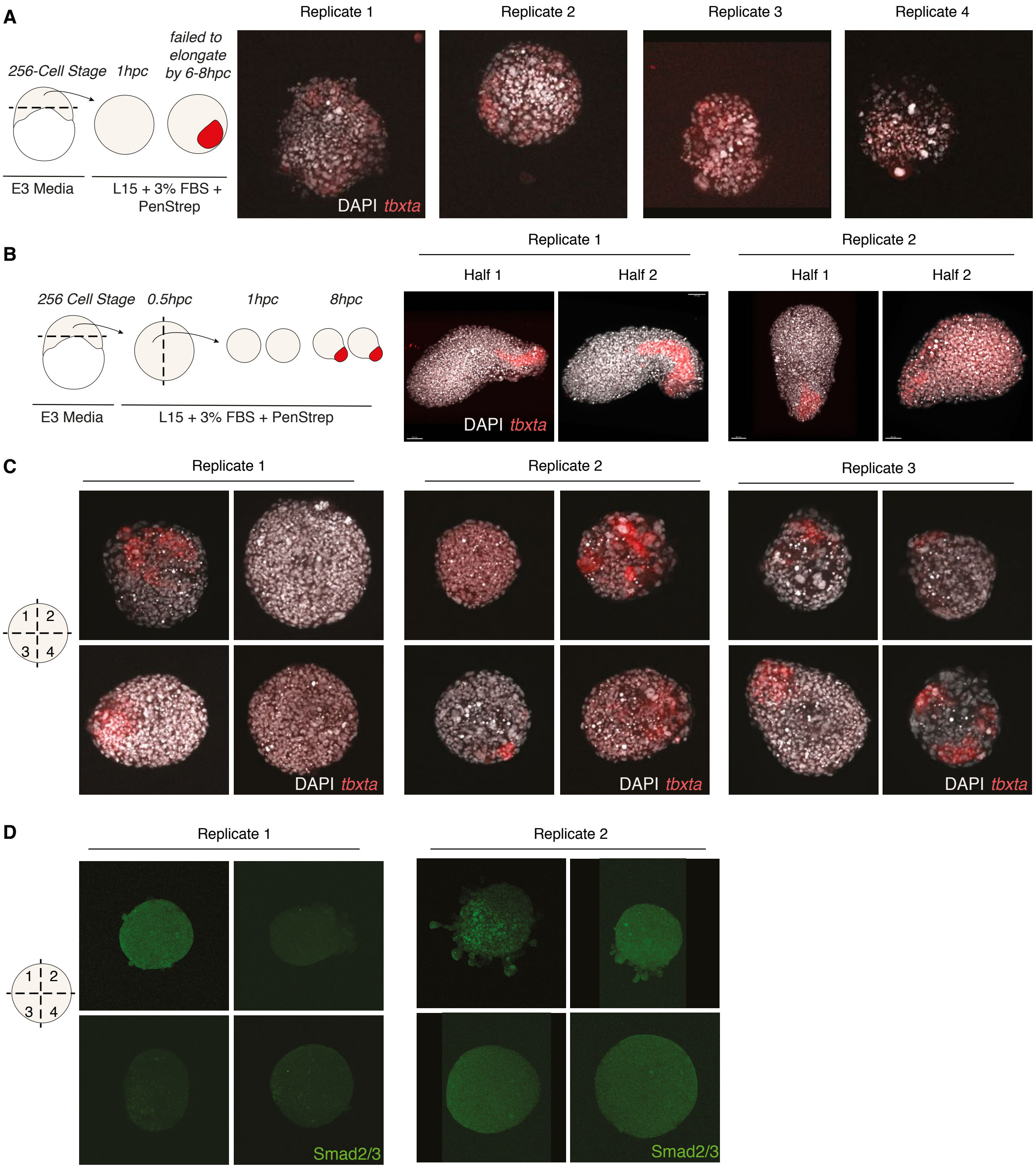

**Figure S2.** Expression of *tbxta* and Smad2/3 activity in explants of the 256 cell stage of different sizes

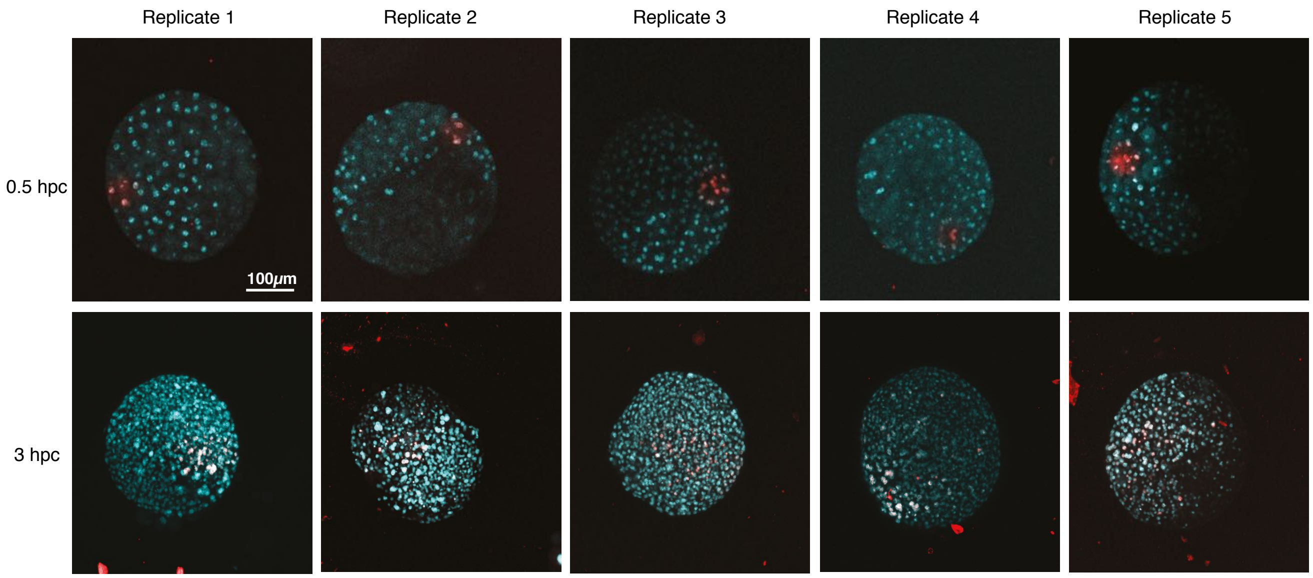

**Figure S3.** Photolabeling experiments

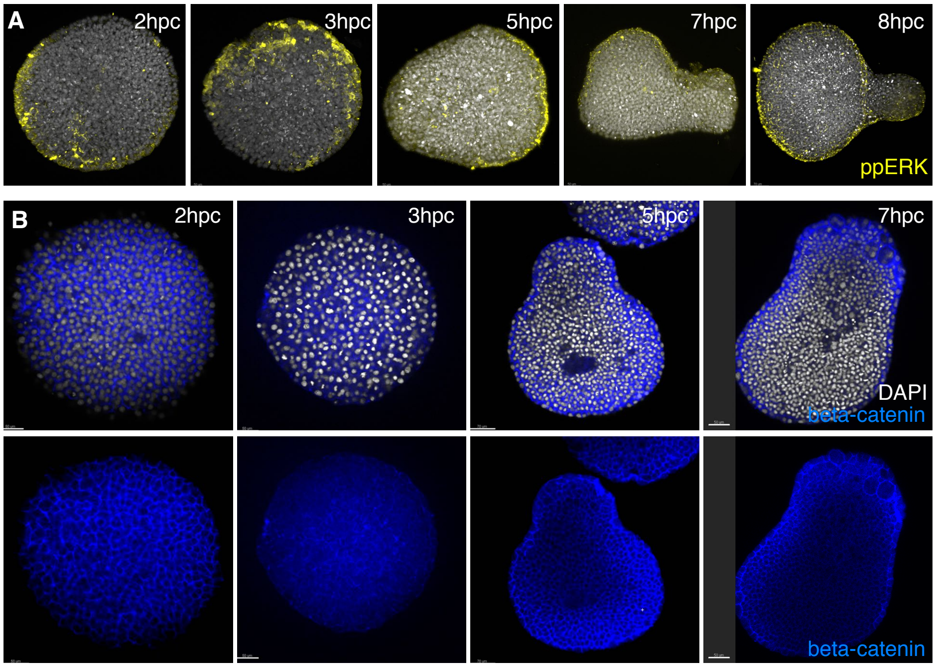

**Figure S4.** FGF, Nodal and BMP activity in pescoids

Effect of different media composition on axial elongation in pescoids

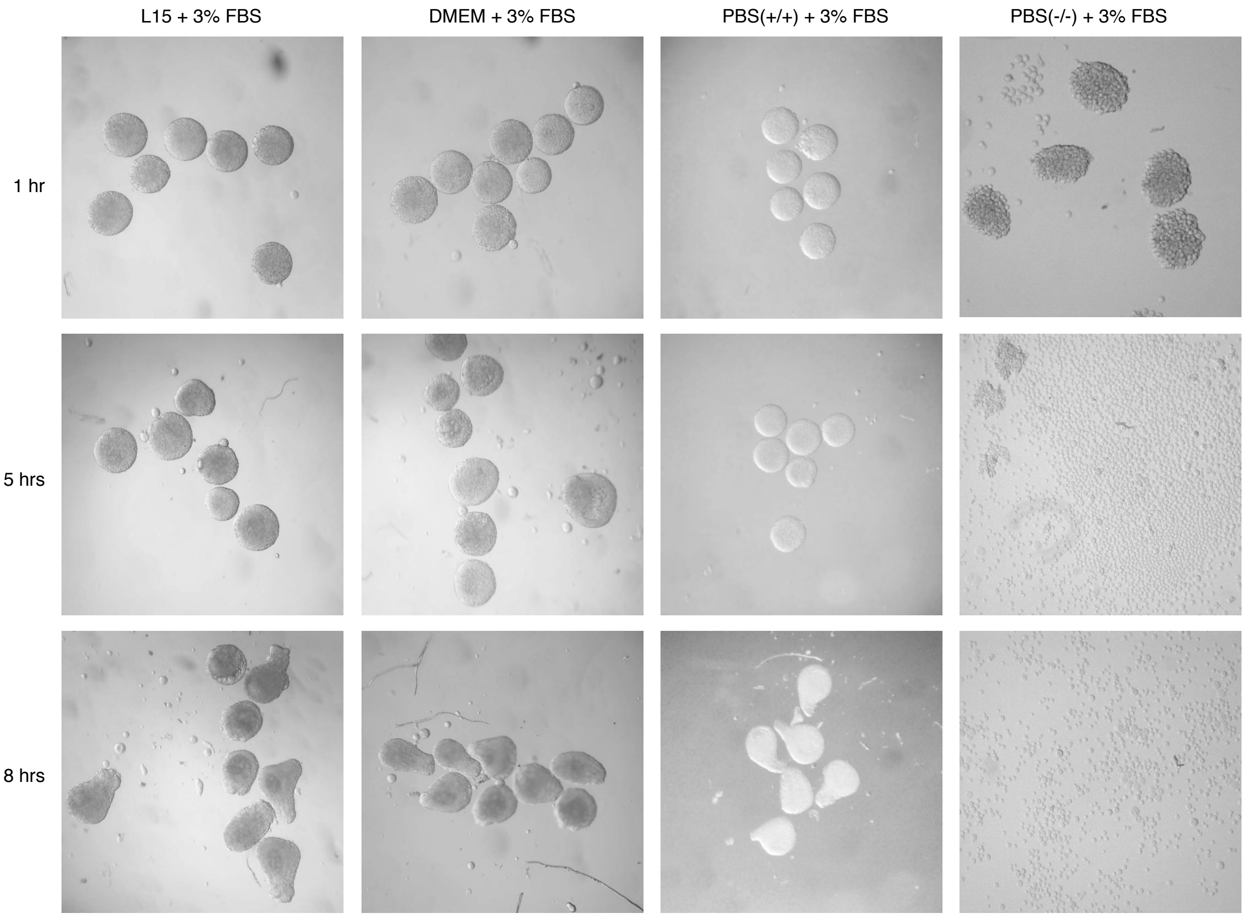

**Figure S5.** Representative images showing the effect of media composition on pescoids. Differences in total number of pescoids between different time points for a given sample type (different media composition) results from pescoids drifting out of the field of view.

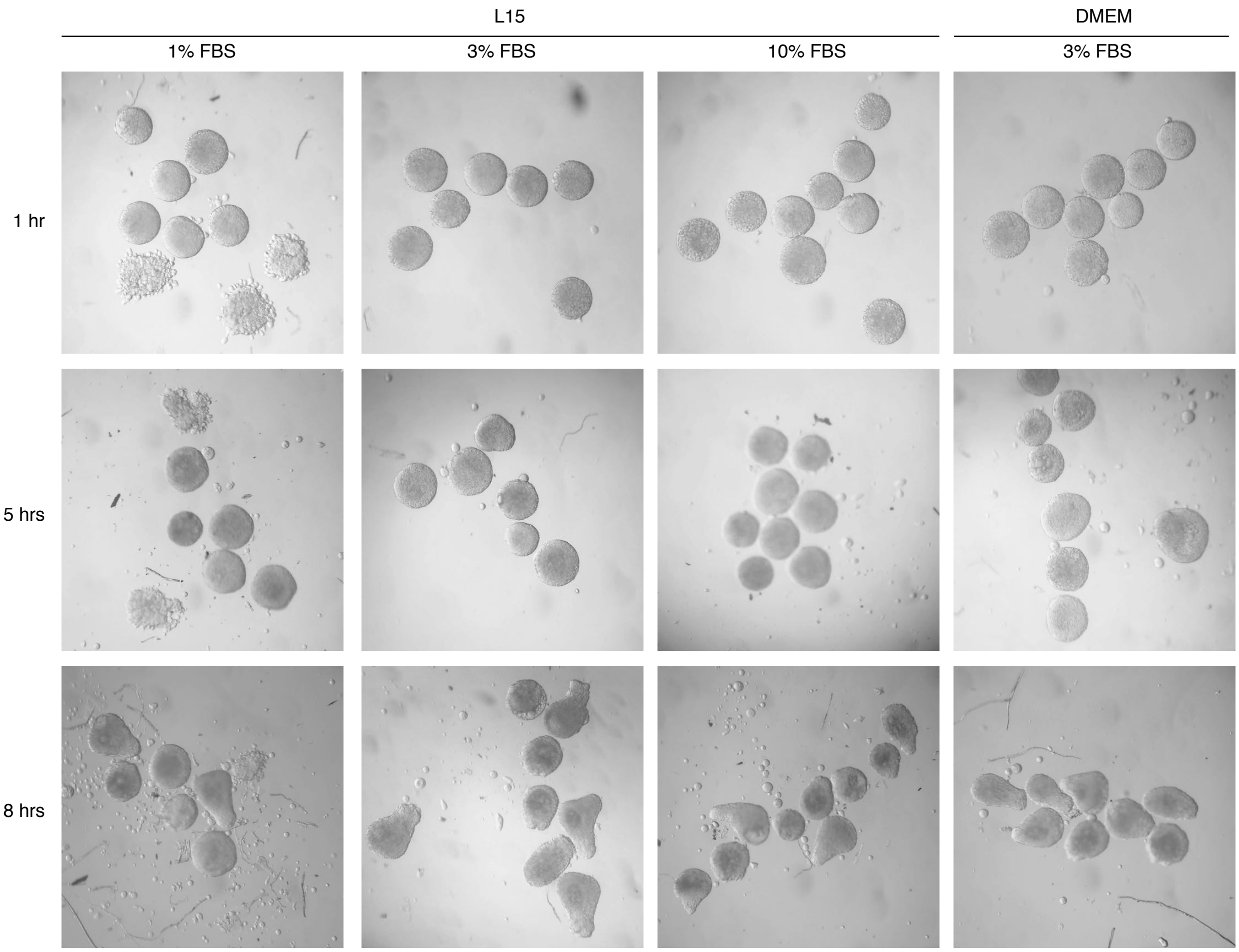

**Figure S6.** Representative images showing the effect of FBS percentage in media on pescoids. Differences in total number of pescoids between different time points for a given sample type (different percentage of FBS) results from pescoids drifting out of the field of view.

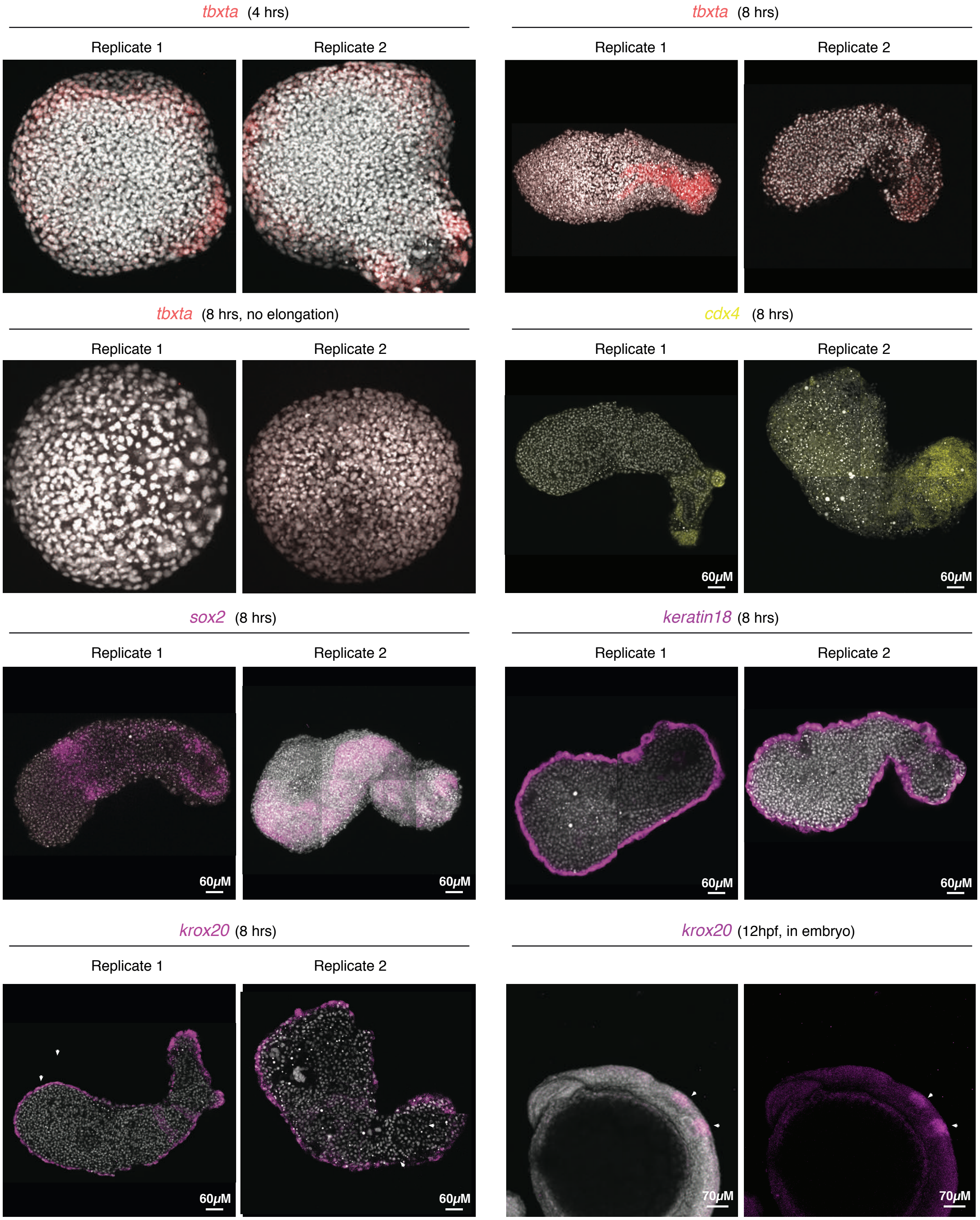

**Figure S7.** Representative images showing HCR stains for *tbxta,krox20,cdx4,sox2* and *keratin18* in elongated pescoids.

### 3 Supplementary Movies

**Video S1** . Brightfield time lapse of an elongating pescoid imaged on a widefield microscope imaged at 10X, one frame per 10 minutes

**Video S2** . Brightfield and fluorescence time lapse of a pescoid containing a Tbx16::GFP reporter imaged at 10X, one frame per 10 minutes.

**Video S3** . Brightfield and fluorescence time lapse of a pescoid containing a TCF::GFP reporter imaged at 10X, one frame per 10 minutes.

**Video S4** . Ubiquitous H2B-GFP labeled pescoids imaged on a SPIM with a time resolution of 2.5 minutes.

**Video S5** . Brightfield and fluorescence time lapse of a single animal cap containing a TCF::GFP reporter imaged at 10X, one frame per 10 minutes.
